# An information-theoretic approach to single cell sequencing analysis

**DOI:** 10.1101/2020.10.01.322255

**Authors:** Michael J. Casey, Jörg Fliege, Rubén J. Sánchez-García, Ben D. MacArthur

**Affiliations:** Mathematical Sciences, University of Southampton, Southampton, UK; Institute for Life Sciences, University of Southampton, Southampton, UK; The Alan Turing Institute, London, UK; Centre for Human Development, Stem Cells and Regeneration, Faculty of Medicine, University of Southampton, Southampton, UK

## Abstract

Single-cell sequencing (sc-Seq) experiments are producing increasingly large data sets. However, large data sets do not necessarily contain large amounts of information. Here, we formally quantify the information obtained from a sc-Seq experiment and show that it corresponds to an intuitive notion of gene expression heterogeneity. We demonstrate a natural relation between our notion of heterogeneity and that of cell type, decomposing heterogeneity into that component attributable to differential expression between cell types (inter-cluster heterogeneity) and that remaining (intra-cluster heterogeneity). We test our definition of heterogeneity as the objective function of a clustering algorithm, and show that it is a useful descriptor for gene expression patterns associated with different cell types. Thus, our definition of gene heterogeneity leads to a biologically meaningful notion of cell type, as groups of cells that are statistically equivalent with respect to their patterns of gene expression. Our measure of heterogeneity, and its decomposition into inter- and intra-cluster, is non-parametric, intrinsic, unbiased, and requires no additional assumptions about expression patterns.

## Introduction

Advances in single-cell sequencing (sc-Seq) technologies have enabled us to profile thousands of cells in a single experiment (Svensson et al. 2018). In combination with advances in unsupervised analysis methods, particularly specialised clustering algorithms and dimensionality reduction techniques, these technologies have allowed us to dissect cellular identities in unprecedented detail and discover novel, functionally important, cell types (Trapnell 2015). The goal of most sc-Seq studies (except those focused on methodology development) is to extract biological information, often concerning the mix of cell types present in the tissue sample, from the data obtained. Yet, data is not the same as information; and large, complex, data sets do not necessarily convey useful or usable information. Notably, current single-cell profiling technologies typically produce noisy data for numerous technical reasons, including low capture rate, sparsity due to shallow sequencing, and batch effects (Kharchenko et al. 2014, Hicks et al. 2018). Consequently, the relationship between biological information and sc-Seq data is complex and incompletely understood. There is, therefore, a need for formal, quantitative methods to evaluate this relationship. To address this challenge, we propose an information-theoretic framework that quantifies the amount of information contained in a sc-Seq data set, and leads to a natural definition of gene expression heterogeneity. Our measure of gene expression heterogeneity decomposes into that which is explained by a given grouping of cells – a proposed clustering into cell types, for example – and that which remains unexplained.

Our framework differs from other approaches to heterogeneity decomposition (e.g. (Lun et al. 2016, Hafemeister & Satija 2019)) by formally quantifying the information, in terms of gene expression heterogeneity, gained from a sc-Seq experiment concerning the mix of cell types present. The resulting measure of heterogeneity, and its decomposition into inter- and intra-cluster heterogeneity, is non-parametric, intrinsic, unbiased, and requires no *a priori* assumptions about gene expression patterns.

Our approach is mathematically precise, biologically intuitive and computationally simple to implement, enabling a practitioner to quickly assess the information content of a sc-Seq data set, in terms of gene expression heterogeneity, identify highly informative genes, and determine the extent to which observed patterns of gene expression variability are explained by the presence of a mixture of cell types in a cellular population. Furthermore, we provide an efficient unsupervised clustering algorithm of cells from sc-Seq data based on heterogeneity and an implementation as an R package.

## Results

High-throughput single-cell analysis methods, such as single-cell sequencing, typically view cells as the objects of study and seek to compare cell identities with each other (Robinson et al. 2010, Brennecke et al. 2013, Grün et al. 2014, Love et al. 2014, Kiselev et al. 2019, Hafemeister & Satija 2019, Townes et al. 2019, Liu et al. 2020). However, this cell-centric view is less well suited to quantifying gene expression heterogeneity, which is concerned with patterns of variation that arise from the mixing of cell types within a population, and may vary from gene to gene. For instance, variance, the standard measure of heterogeneity in the cell-centric view, may not be informative in multi-modal distributions (Smith & MacArthur 2017).

In the context of sc-Seq, heterogeneity results from differential expression between cell types. In the simplest case, where one cell type expresses a gene highly and another lowly, such differential expression introduces bimodality in the cell-centric expression distribution. In a bimodal distribution, the population mean is no longer characteristic of either of the two subpopulations, making variance, as a measure of the spread about the mean, also misleading (see Smith & MacArthur (2017)).

To meet these challenges, we introduce a novel gene-centric probabilistic view that seeks to more formally specify what is meant by gene expression homogeneity and heterogeneity. We show that this gene-centric view is better suited to quantifying gene expression heterogeneity in the context of a multimodal population, and formalises the biological intuition of expression homogeneity and heterogeneity.

### Quantifying gene expression heterogeneity

Consider the expression pattern of a single gene *g* of interest in a population of *N* distinct cells. Assume that in total *M* transcripts of *g* are identified in the cell population (i.e. across all *N* cells profiled). Note that *M* represents the observed transcript count, which may differ from the true count due to technical artefacts. Now consider the stochastic process of assigning the *M* identified transcripts of gene *g* to the *N* cells profiled. Intuitively, the population is homogeneous with respect to expression of *g* if all the cells are statistically the same with respect to its expression. Mathematically, this means that the *M* transcripts of *g* will be assigned to the *N* cells independently and equiprobably – i.e. each transcript will be assigned to each cell with probability 1*/N*. Conversely, if the population is heterogeneous with respect to expression of *g* (that is, it consists of a mix of cell types, each expressing the gene differently), then transcripts will not be assigned uniformly, but rather will be assigned preferentially to distinct subsets of cells. Heterogeneity in experimentally observed patterns of expression can, therefore, be assessed in terms of deviation from this hypothetical homogeneous null model.

The Kullback-Leibler divergence (KLD, also known as the relative entropy) is a measure of the information encoded in the difference between an observed and null distribution (Kullback & Leibler 1951). In the sc-Seq context, the KLD of an experimentally observed gene expression distribution from the homogeneous null model, described above, measures the amount of information that is lost by assuming that the gene is homogeneously expressed in the sequenced cell population. We will denote this quantity *I*(*g*) and refer to it as the *heterogeneity* of *g* (see **Fig. 1** for a schematic). Formally,

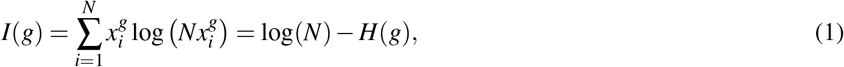

where 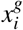 is the fraction of transcripts of gene *g* expressed by cell *i*, for each 1 *≤ i ≤ N*, and *H*(*g*) is the entropy of the expression of *g* in the population (see **Methods** for full details). For technical reasons, explained in the **Methods**, we will use natural logarithms in all calculations; *I*(*g*) therefore has units of nats.

**Figure 1.**
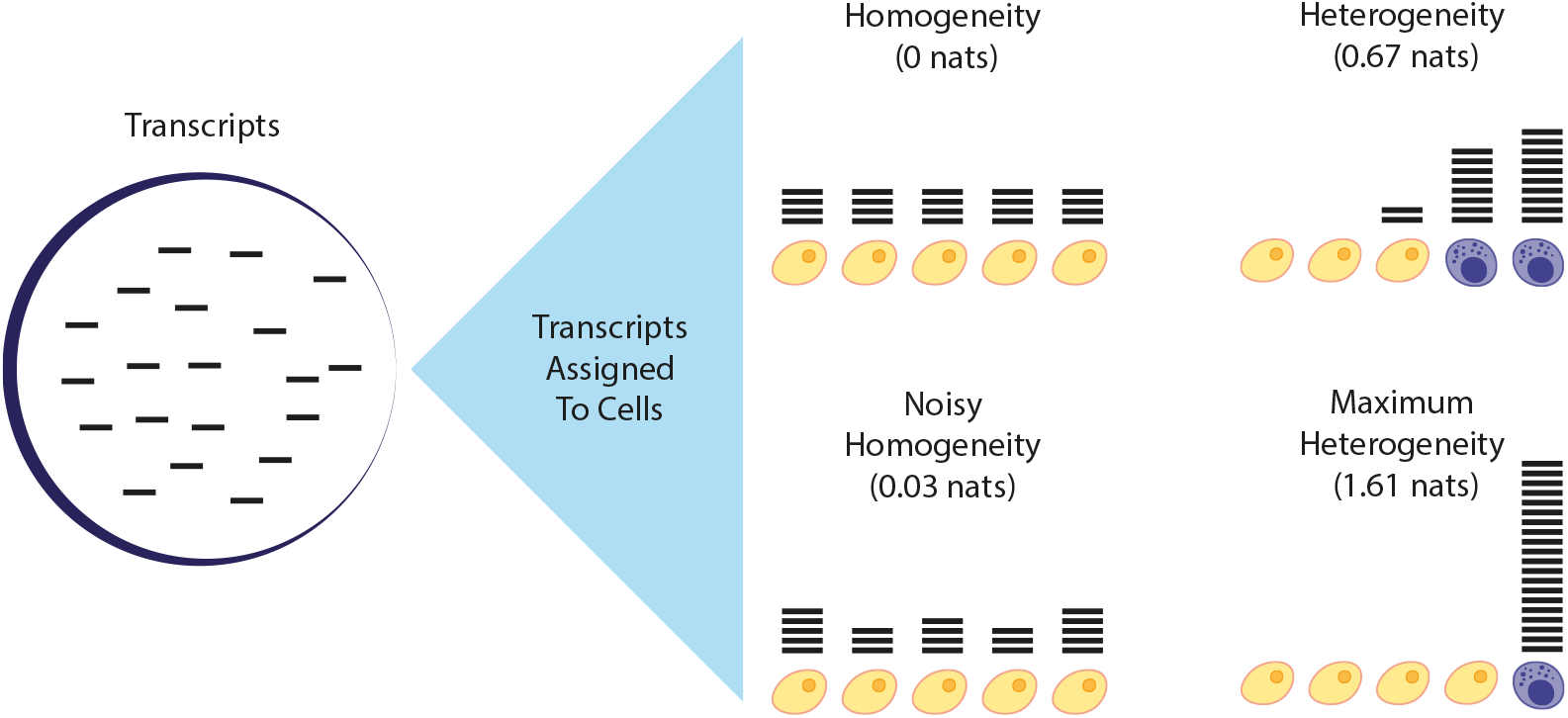
An information-theoretic view of sc-Seq data. Transcripts, or more generally counts, of a given gene (shown here as horizontal bars) are assigned to cells after sequencing. If the cell population is homogeneous with respect to the expression of *g*, then the heterogeneity *I*(*g*) will be zero (top left population, *I*(*g*) = 0). In practice, the transcript assignment process is stochastic, and so there will always be some deviation from this ideal (bottom left population, *I*(*g*) small). (Note that the technical effects of this stochasticity on the information obtained may be reduced by using a shrinkage estimator to determine the distribution of transcripts (see **Methods**)). If the population is heterogeneous, then transcripts may be preferentially expressed in a subset of cells and the information obtained from the experiment, as measured by *I*(*g*) will be larger (top right population, *I*(*g*) large), reaching a maximum at log(*N*), where *N* is the number of cells sequenced, when only one cell expresses the gene (bottom right population, *I*(*g*) = ln(5) *≈* 1.61 largest). Note that the population heterogeneity *I*(*g*) is independent of any decomposition of the cell populations into subpopulations (shown here as yellow and purple cells, for illustration). However, given any grouping of the cells into subpopulations, *I*(*g*) can be formally decomposed as the sum of the heterogeneity explained by within and in-between subpopulations (see **Results** and **Fig. 3**). This decomposition, but not the overall value of *I*(*g*), does depend on the chosen assignment of cells to subpopulations.

Intuitively, if the cell population is unstructured with respect to the expression of *g* (i.e. if the cells are approximately interchangeable with respect to the expression of *g*) then the assumption of homogeneity is correct and *I*(*g*) *≈* 0. Conversely, the theoretical maximum for *I*(*g*) is log(*N*), which is achieved when *H*(*g*) = 0 and all transcripts of the gene are assigned to the same cell (see **Fig. 1**). Note that: (1) we do not need to know, or model, the particular expression distribution of *g* in the population, so no *a priori* assumptions about expression patterns are required to calculate *I*(*g*); (2) *I*(*g*) is agnostic concerning missing readings or counts so long as they are distributed uniformly at random; (3) since it quantifies the deviation from the homogeneous null model, *I*(*g*) measures the information obtained from the experiment concerning the expression of *g*.

In practice, *I*(*g*) is associated with cellular diversity: the more distinct cell sub-populations present in a sample, and the more those sub-populations differ from one another with respect to their expression of *g*, the larger *I*(*g*) will be. Thus, *I*(*g*) is a parsimonious measure of expression heterogeneity that makes minimal assumptions concerning expression patterns and, therefore, imposes minimal technical requirements on data collection methodology or quality. As such, it can be used as the basis for numerous aspects of the sc-Seq pipeline, including feature selection and unsupervised clustering.

To validate these uses, we considered a series of single-cell RNA-sequencing bench-marking data sets: *Svensson*, a technical control (Svensson et al. 2017); *Tian*, a mixture of three cancerous cell lines (Tian et al. 2019); *Zheng*, a set of FACS separated peripheral blood mononuclear cells (PMBCs); *Stumpf*, a sampling of cells from mouse bone marrow (Stumpf et al. 2020); and the *Tabula Muris*, a mouse cell atlas with cells from twelve organs (Tabula Muris Consortium 2018). **Fig. 2a-e** shows plots of *I*(*g*) for each gene profiled in these experiments. These plots confirm the intuition that the more distinct types present in a sample, and the more those cell types differ from one another with respect to their gene expression patterns, the greater the population heterogeneity will be, as measured by *I*(*g*).

**Figure 2.**
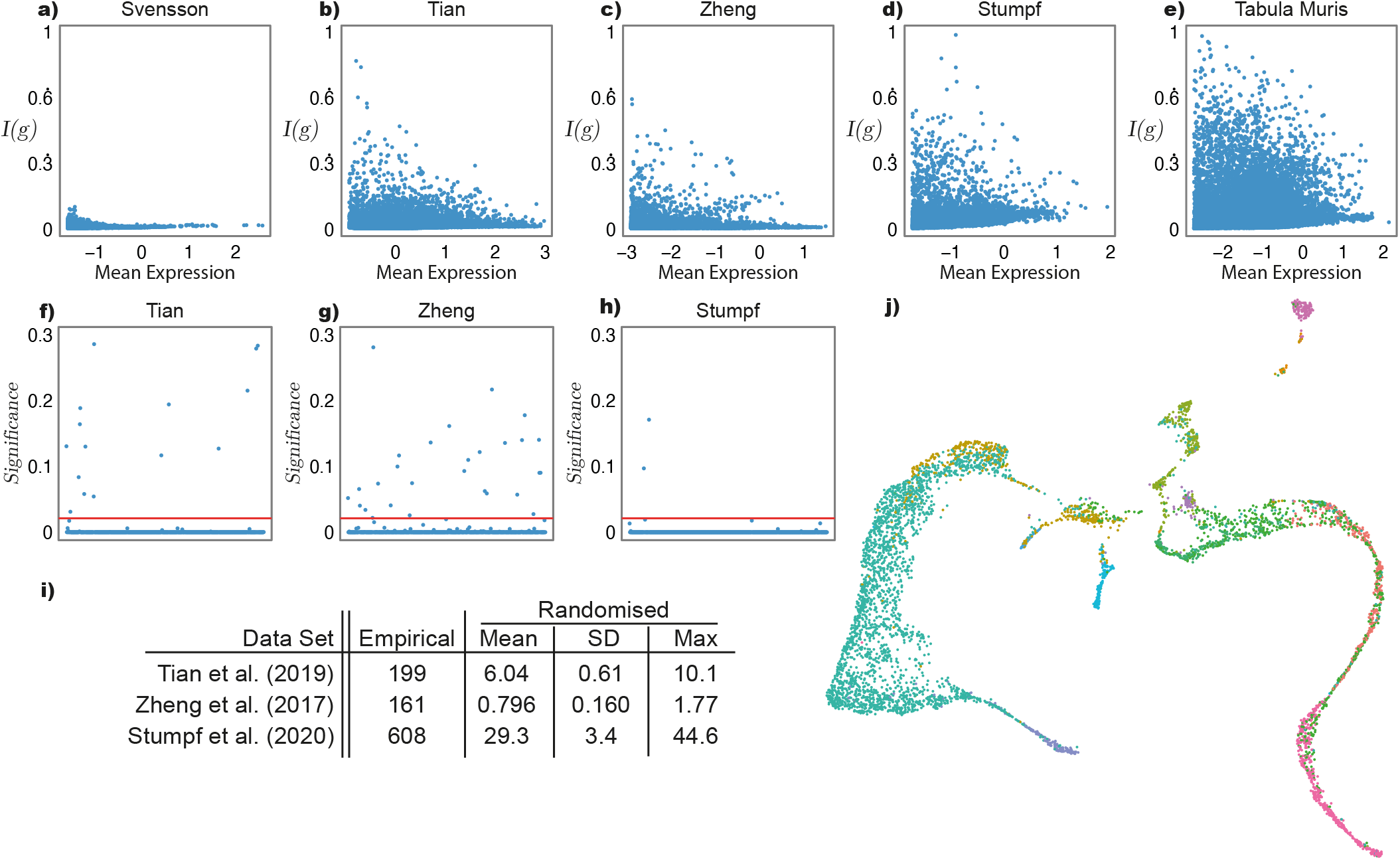
Information-theoretic single-cell analysis. Recall that *I*(*g*) measures the heterogeneity of a cellular population with respect to the expression of *g*: *I*(*g*) = 0 when transcripts are expressed uniformly and increases as transcripts are expressed preferentially in a subset of cells, reaching a maximum *I*(*g*) = log(*N*), where *N* is the number of cells sequenced, when only one cell expresses the gene. **a-e)** Plots of expression heterogeneity, *I*(*g*) (normalised by the theoretical maximum, log(*N*)) against log mean expression for the bench-marking sc-Seq data sets described in the main text. In each panel, each point represents a gene profiled. The number of genes associated with large values of *I*(*g*) increases with the number of cell types present in the population profiled, showing *I*(*g*) as a valid measure of cell type diversity. Panel **a)** shows data from a technical control (Svensson et al. 2017) (number of cell types, *C* = 1), **b)** a mixture of three cancerous cell lines (Tian et al. 2019) (*C* = 3), **c)** FACS sorted immune cells (Zheng et al. 2017) (*C* = 4), **d)** a sample of mouse bone marrow (Stumpf et al. 2020) (*C* = 14), and **e)** a multi-organ mouse cell atlas (Tabula Muris Consortium 2018) (*C* = 56). **f-h)** Biologically meaningful cell annotations are associated with high inter-cluster heterogeneity. Established cell annotations for the **f)** *Tian*, **g)** *Zheng* and **h)** *Stumpf* data are associated with higher inter-cluster heterogeneity than expected by chance (i.e., in randomly permuted clusters; significance is assessed using a one-sided exact test with 10^4^ permutations; y axes show log_10_(*p* + 1)). In all panels the red line shows *p <* 0.05, false discovery rate corrected for 500 trials (Benjamini & Hochberg 1995, Fisher 2017). Genes below this threshold are significantly different gene expression patterns across the set of identified cell types. **i)** Summary statistics for the total inter-cluster heterogeneity *H*_*S*_ = ∑_*g*_ *H*_*S*_(*g*) based on established empirical and randomly permuted cell annotations (10^4^ random permutations in each case). These statistics show the strong association of high *H*_*S*_ with biologically meaningful groupings of cells. **j)** A Uniform Manifold Approximation and Projection (UMAP) (McInnes et al. 2018) plot of the top 500 genes by *I*(*g*) for the *Stumpf* data set; each point is a cell, coloured by its scEC cluster. This shows that *I*(*g*) is able to capture the continuous variation of developing cell types.

These results indicate that heterogeneity (as defined here) can be used for feature selection. As a measure for use in feature selection, *I*(*g*) identifies gene sets largely distinct from those based on Highly Variable Gene selection (the mean overlap of selected genes – for listed data sets excluding *Svensson* – was 0.36, with the number of genes selected determined by scran, based on an false discovery rate threshold of 0.05). For instance, *I*(*g*) identifies *Hbb-bs*, a marker of erythrocyte maturation, as informative in the Stumpf et al. (2020) data set (high *I*(*g*) value), whereas scran does not, instead attributing the observed variance to technical sources (see **Supplementary Figure 1**)(Lun et al. 2016). Critically, information-theoretic heterogeneity is more mathematically precise and computationally simpler to implement than variance-based approaches, making no distributional assumptions (e.g. use of the negative binomial model) and having no free parameters to fit. Furthermore, it may be easily modified to account for the presence of multiple homogeneous cell subpopulations, and thereby act as a measure of cluster quality.

### Inter-cluster Heterogeneity

Let *S* be a discrete clustering of a population of cells – i.e. an assignment of the cells to a finite set of *C* non-intersecting sub-populations (also known as clusters). In the **Methods**, we show that *I*(*g*) can be decomposed into two parts: one part that quantifies the extent to which each of the sub-populations defined by *S* deviate from homogeneity, which we call the *intra-cluster heterogeneity* and write as *h*_*S*_(*g*); and one part that quantifies how differently the gene is expressed, in aggregate, between sub-populations, which we call the *inter-cluster heterogeneity* and write as *H*_*S*_(*g*). Namely, we show that

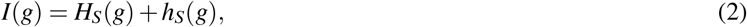

for any proposed clustering *S*. Thus, the information obtained from an experiment concerning the expression of a gene *g* can be explicitly related to both local (within cluster) and global (between cluster) patterns of variation, for any proposed clustering. Full mathematical details of this decomposition, including formulae for *H*_*S*_(*g*) and *h*_*S*_(*g*), are provided in the **Methods**, and an illustration is given in **Fig. 3**. We also provide an R package for its calculation (see **Methods**).

**Figure 3.**
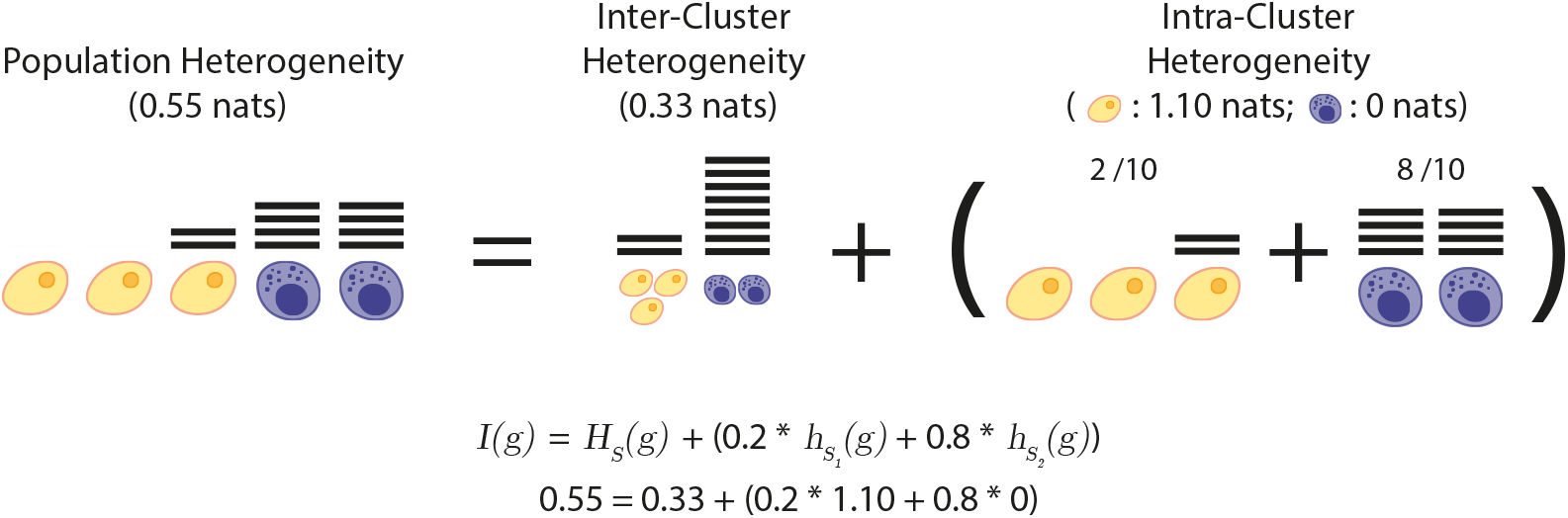
Heterogeneity is additively decomposable. The heterogeneity of a population of cells (5 cells in this illustration) with respect to the expression of a gene *g, I*(*g*), can be decomposed into inter- and intra-cluster heterogeneities for any proposed clustering, *S* (here, two subpopulations, or clusters, of 3 yellow and 2 purple cells). The inter-cluster heterogeneity *H*_*S*_(*g*) is determined by independently aggregating all transcripts (shown as horizontal lines) associated with each sub-population in *S* and then taking the KLD of the resulting distribution from the uniform distribution of the transcripts over *C* clusters. It measures the extent to which transcripts are uniformly assigned to clusters. The intra-cluster heterogeneity *h*_*S*_(*g*) is determined by taking the weighted sum (with respect to the number of transcripts on each subpopulation) of the heterogeneities of each of the constituent subpopulations, considered independently. It represents the average heterogeneity of the proposed clusters, accounting for disparities in number of transcripts assigned. In this toy example, the overall population heterogeneity of gene *g, I*(*g*) = 0.55, decomposes as the sum of the inter-cluster heterogeneity *H*_*S*_(*g*) = 0.33, plus the intra-cluster heterogeneity *h*_*S*_(*g*) = 0.22. The latter is obtained as the weighted sum (with respect to the number of transcripts in each cluster, here 2*/*10 = 0.2 and 8*/*10 = 0.8) of the heterogeneities on each subpopulation. Further details and formulae are provided in the **Methods**.

This decomposition can be used to assess cluster quality. For a proposed cluster to meaningfully represent a cell type, it must be associated with differential expression of some subset of marker genes. Because *H*_*S*_(*g*) measures the extent to which the expression of gene *g* deviates from the homogeneous null model it is a simple measure of the extent to which *g* is differentially expressed between the clusters defined by *S*. We therefore expect that biologically meaningful clustering based on a marker gene *g* will result in a high value for *H*_*S*_(*g*) and (by necessity) a low value for *h*_*S*_(*g*).

This expectation is confirmed for the *Tian, Zheng* and *Stumpf* data sets for which we possess high-confidence *a priori* cell type annotations, derived either experimentally (for the *Tian* and *Zheng* data) or from expert annotation of a computational clustering (for the *Stumpf* data). In the set of genes most likely to be differentially expressed between clusters in these data sets (taken to be the top 500 genes by *I*(*g*) in each case) the majority are associated with substantially higher inter-cluster heterogeneity than expected by chance (i.e., in randomly permuted clusters). Indeed, in *Tian* 97.0%, in *Zheng* 94.4% and in *Stumpf* 99.6% of the tested genes had significantly higher inter-cluster heterogeneity than expected (one-sided exact test with 10^4^ permutations, *p <* 0.05, false discovery rate corrected for 500 trials; see **Fig. 2f-h** for illustration) (Benjamini & Hochberg 1995, Fisher 2017).

These results indicate that by simple information-theoretic reasoning, we are able to quantify the amount of information in the expression pattern of a single gene that is explained by a given clustering. However, a key strength of high-throughput single-cell profiling methods is that they allow the simultaneous profiling of thousands of genes in large cell populations.

Our information-theoretic reasoning may be extended to a high-throughput single-cell profiling experiment by assuming that each gene is an independent source of information and making use of the fact that information from independent sources is additive (Shannon 1948). Thus, we can determine the total information explained by a given clustering *S* by evaluating the sum *H*_*S*_ = ∑_*g*_ *H*_*S*_(*g*) over all genes profiled. The total information explained by *S* is a simple, easily computed measure for cluster evaluation that favours grouping of cells into homogeneous (with respect to gene expression) sub-populations and is maximised at ∑_*g*_ *I*(*g*) if and only if the proposed clustering divides the population into indivisible perfectly homogeneous sub-populations and thereby accounts for all the heterogeneity contained in the sc-Seq data set. If so, then *S* is the maximum entropy partition of the cell population into distinct classes, and may therefore be considered as the most parsimonious way of assigning cell identities.

To illustrate the association between total information explained, *H*_*S*_, and cluster quality, we again considered the established annotations of the *Tian, Zheng* and *Stumpf* data sets. As expected, in all cases the observed value of *H*_*S*_ significantly exceeds that of all random label permutations (see **Fig. 2i**) indicating that *H*_*S*_ is strongly associated with cluster quality and that these benchmark cell annotations are associated with homogeneous cell sub-populations. Notably, this association holds true independently of the method used to annotate cell identities (annotations were derived from genotypic information for the *Tian* data, surface protein expression for the *Zheng* data, and unsupervised clustering for the *Stumpf* data), indicating that *H*_*S*_ provides a methodology agnostic, parameter-free, means to quantify cluster quality.

### Unsupervised Clustering

Building on this empirical association, we investigate the clustering of cells that maximises *H*_*S*_ (and necessarily minimises *h*_*S*_) as a reasonable approximation to (homogeneous) cell type. That is, this is the cell clustering that explains the most of the gene expression heterogeneity as inter-cluster variability.

We implement *H*_*S*_-maximisation as the objective function of an unsupervised clustering method, which we call scEC (single-cell Entropic Clustering). We show that our method is comparable to the state-of-the-art Louvain method, and in doing so, validate the underlying definition of heterogeneity and homogeneous cell type.

Namely, we compare the clustering returned by scEC and the established annotations for the *Tian, Zheng* and *Stumpf* data sets. The resulting scEC clusterings agreed strongly with the established cell annotations, achieving adjusted Rand Indices (ARI) of 0.99 for the *Tian* annotations, 0.87 for the *Zheng* annotations, and 0.69 for the *Stumpf* annotations (the adjusted Rand index is a measure of similarity of a proposed clustering to a known clustering, Rand (1971)). To benchmark these results we also repeated the analysis using the Louvain method, a leading single-cell clustering algorithm, which achieved ARIs of 0.99, 0.99 and 0.35 for the *Tian, Zheng*, and *Stumpf* data sets respectively (Blondel et al. 2008, Luecken & Theis 2019, Freytag et al. 2018, Stuart et al. 2019).

The scEC objective function, *H*_*S*_, is naturally built by summation of gene-level measurements (see **Methods**), so can be used to directly identify the key drivers of a given clustering without relying on *post-hoc* differential gene expression testing. To demonstrate this feature, we compared the inter-cluster heterogeneity values (i.e., the values *H*_*S*_(*g*) for each of the genes profiled) for clusters identified by scEC and by established annotations for the *Tian & Stumpf* data sets (see **Fig. 4**).

**Figure 4.**
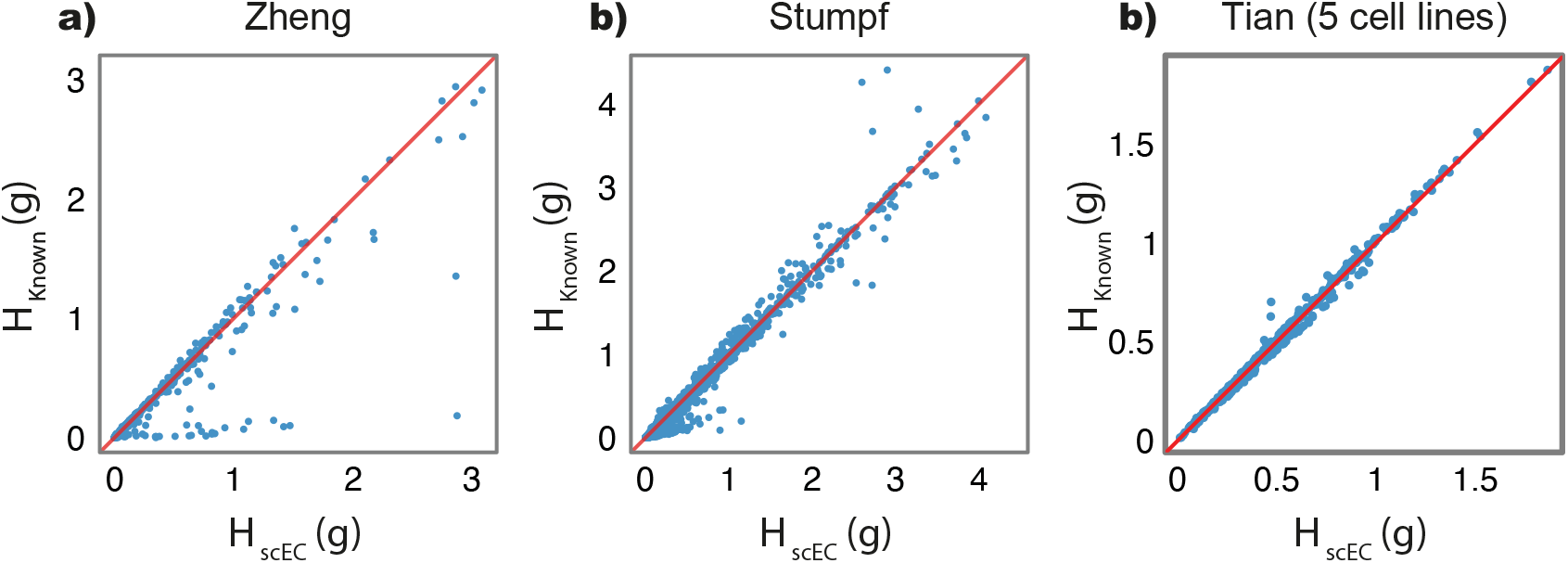
Comparison of inter-cluster heterogeneity of scEC-generated clusters versus established annotations. Plots of *H*_*S*_(*g*) based on an scEC-generated clustering (*x*-axis) and established annotations (*y*-axis) for the **a)** *Zheng* and **b)** *Stumpf*, and **c** an alternative data set from *Tian* (Zheng et al. 2017, Stumpf et al. 2020, Tian et al. 2019). In all panels each point represents a gene profiled and the red line indicates *H*_*scEC*_(*g*) = *H*_*Known*_(*g*). For the genes below the red lines, the scEC clustering is better than the prior annotation at explaining the gene expression heterogeneity as inter-cluster variability, and vice versa for the genes above the red line.

We found that the values of *H*_*S*_(*g*) associated with scEC clusters are strongly positively correlated with those associated with established annotations for both the *Zheng* and *Stumpf* data sets (Pearson’s correlation coefficient of 0.94 for *Zheng* and 0.99 for *Stumpf* ; see **Fig**. 4). Moreover, the key marker genes in each data set were found to have significantly different expression levels across clusters, as measured by *H*_*S*_(*g*) (one-sided permutation test on clustering labels, 10^4^ permutations, *p <* 0.05, false discovery rate corrected; marker genes for *Zheng* : *CD14, CD4, CD8* and *NCAM1*, Zheng et al. (2017); and *Stumpf* : *Cd34, Kitl, Spi1, Gata1* and *Pax5*, Stumpf et al. (2020)).

This concordance indicates that scEC is generally able to recapitulate known cell clusters in an unbiased and parameter free way. However, despite this concordance, there were differences between scEC clusters and established annotations. To investigate these differences further we selected those genes for which the value of *H*_*S*_(*g*) differed substantially (Δ*H*_*S*_(*g*) = |*H*_*scEC*_(*g*) *−H*_*Known*_(*g*)| *>* 0.1 nats) between scEC clusters and established annotations in each data set for further investigation. In the *Zheng* data, we found that those genes with expression heterogeneity better explained by the scEC clustering were enriched for cell-cycle related genes (gene ontology enrichment, *p* = 1.33 *×* 10^−8^, false discovery rate corrected). Conversely, those genes with expression heterogeneity better explained by the established annotation were enriched for genes involved in the immune response (*p* = 1.49 *×* 10^−3^, false discovery rate corrected) (Ashburner et al. 2000, Gene Ontology Consortium 2021, Benjamini & Hochberg 1995).

In the *Stumpf* data set, we found that those genes with expression heterogeneity better explained by the scEC clustering were enriched for genes involved in erythroblast differentiation and homeostasis (*p* = 1.80 *×* 10^−2^ and *p*-value = 6.17 *×* 10^−4^ respectively, false discovery rate corrected). Conversely, those genes with expression heterogeneity better explained by the established annotation were enriched for genes involved in the immune response (*p* = 2.57 *×* 10^−18^, false discovery rate corrected). In this example, scEC generally identifies additional erythrocyte sub-populations while merging the cell sub-types of the neutrophil lineage (see **Fig. 2j** for illustration and **Supplementary Table** for contingency table of cluster assignments). Collectively, these differences reflect the preference of scEC for cellular annotations driven by biological processes involving larger tranches of genes. Depending on circumstance, this preference may be a benefit or a drawback. It is generally beneficial because it ensures that scEC annotations are robust to outliers (i.e. anomalous expression patterns of individual genes cannot easily distort scEC cluster assignments). Conversely, in circumstances in which key cellular functions are determined by a small number of genes, scEC may fail to disambiguate important cell populations from similar cell types. However, given the simplicity of the scEC methodology this issue can this issue can be relatively easily addressed, by adjusting the prior, as follows.

In the standard formulation of scEC each gene contributes equally to the objective function *H*_*S*_ = ∑_*g*_ *H*_*S*_(*g*). This corresponds to assuming a uniform prior. However, alternatives may be easily considered that take into account different sources of prior knowledge. In general, each gene can contribute in a weighted way to the overall inter-cluster heterogeneity. In this case the objective becomes *H*_*S*_ = ∑_*g*_ *w*(*g*)*H*_*S*_(*g*), where *w*(*g*) is the weight of gene *g*. These weights can encode known biology – for example, by up-weighting known drivers of cellular identities, down-weighting genes associated with housekeeping processes and/or the cell cycle, or by weighting genes in inverse proportion to the size (i.e. the number of associated genes) of their leading ontology term.

While incorporating such priors into scEC may be beneficial, defining them is a challenge and potentially introduces a source of bias into the resulting clustering and so should be approached with care. An alternative approach – which encodes prior information in a data-led, rather than algorithmic way – is to use a reference data set, such as a cell atlas, to benchmark cell annotations (these atlases can serve as *reference transcriptomes*, in analogy to reference genomes).

Doing so leads to a semi-supervised version of scEC, wherein we cluster a novel test data set based on the known clustering of a given reference data set. We detail the mathematics of semi-supervised clustering in the **Methods**, but, in short, the extension to the semi-supervised setting requires no additional mathematical machinery. Such mathematical simplicity affirms information theory as providing a mathematically precise and biological intuitive framework for cellular clustering.

As an example, we cluster the *Tian* data set based on a distinct data set containing the same three cell lines (plus two additional cell lines) (Tian et al. 2019). The semi-supervised clustering strongly agreed with the established cell annotation, achieving an ARI of 0.90. Notably, the ARI was reduced compared with the unsupervised version due to the assignment of *∼* 6% of cells to cell types present only in the reference data set, a disadvantage of semi-supervised (and supervised) methods.

### Benchmarking & ImputationC

We benchmark scEC and state-of-the-art methods against a series of data sets consisting of cells sampled from three or five distinct cancerous cell lines and sequenced using various technologies (CEL-Seq2, Sort-seq, 10x, Drop-seq) (Tian et al. 2019, Hashimshony et al. 2016, Peterman & Levine 2016, Macosko et al. 2015). We compare each method’s clustering against ground truth for each data set (by adjusted rand index), finding that scEC performs subpar to the best of existing methods (Grün et al. 2015, Lun et al. 2016, Kiselev et al. 2017, Risso et al. 2018, Stuart et al. 2019). Investigating the gene-wise contributions via *H*_*S*_(*g*) reveals that scEC-derived clusterings diverge from the ground truth due to genes expressed in small numbers of cells, e.g., SPINK6 (difference in *H*_*scEC*_ from ground truth of 0.23 nats; expressed in 1.6% of cells in the five-cell line, 10X data set; see **Fig. 4c**).

To protect against low cell-count genes, we adopt a data-diffusion step, imputing the expression value of each cell as a weighted average of itself and a small subset of cells with similar expression patterns (i.e., diffusion of observed expression values over a shared nearest-neighbours graph derived from the top 500 genes by *I*(*g*), see **Methods**) (Van Dijk et al. 2018). The smoothing layer shares information between genes, lessening scEC’s sensitivity to the expression pattern of individual genes; this raises the performance of scEC clustering to be on par with the state-of-the-art (see **Fig. 5a**).

**Figure 5.**
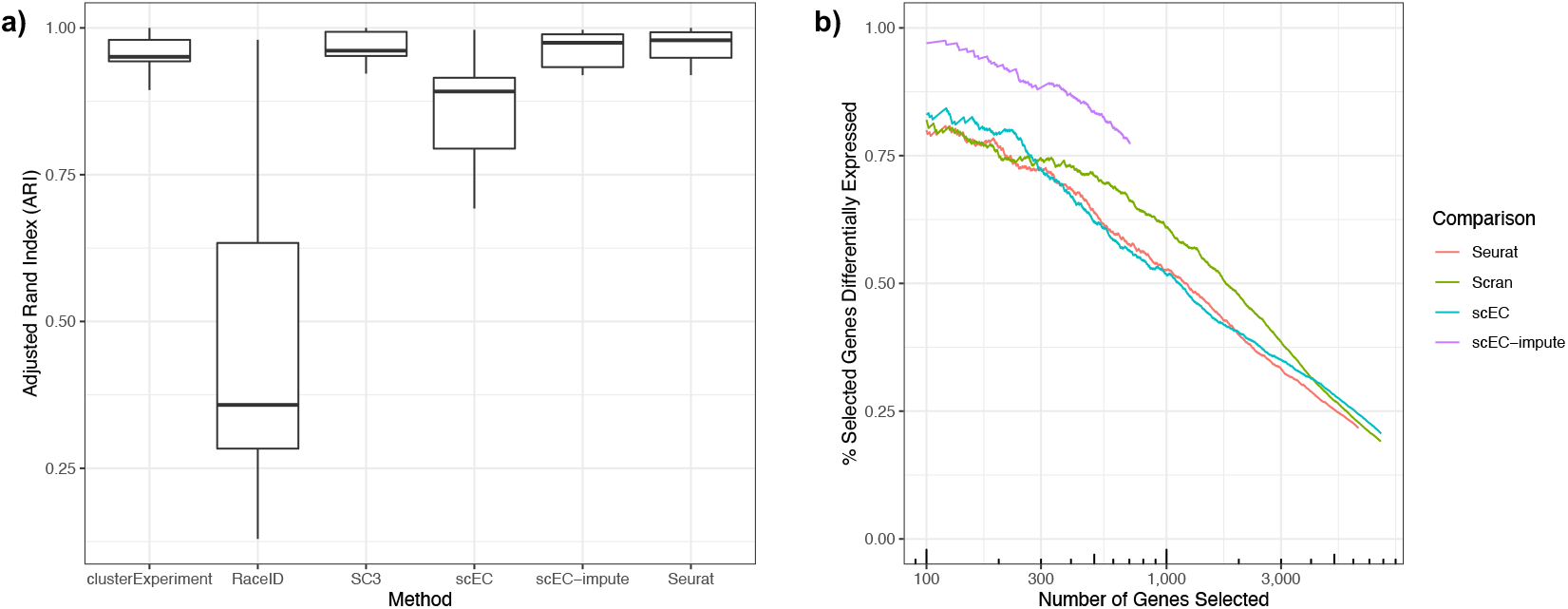
Benchmarking of scEC performance in unsupervised clustering and feature selection. **a)** Adjusted Rand Index of clusterings produced by specified methods against known ground truth for seven data sets, each consisting of three or five cancerous lines sequenced on different platforms. With an additional imputation step, scEC performs on par with other methods. **b)** The percent of the top *N* genes by different feature selection metrics that are differentially expressed. Data set is Sc-seq from three cancerous cell lines sequenced by Drop-seq (with 2005 differential expressed genes identified from non-parametric testing for each cell line versus the remaining; Wilcox test, fasle discovery rate corrected *p*-value *<* 0.05). The greater ability of scEC-impute to *a priori* select differentially expressed genes is repeated across each benchmark data set, see **Supplementary Figure 2**. Note that the imputation step in scEC-impute assigns many genes a heterogeneity *I*(*g*) of zero, resulting in a low cut-off on total number of selectable genes.

We further benchmark scEC (with and without the imputation step) as a feature selection tool, evaluating the ability of *I*(*g*) to *a priori* identify differentially expressed genes, assuming differential expression to be the ground truth of feature selection (differentially expressed genes are identified per data set via Wilcoxon Rank Sum testing, selecting those genes with an FDR-adjusted *p*-value *<* 0.05 in at least one cluster; Stuart et al. (2019)). We find that, without smoothing, scEC is comparable to the state-of-the-art; with smoothing, scEC notably improves, substantially bettering the state-of-the-art in selecting differentially expressed genes from the benchmark data sets (see **Fig. 5b**). Notably, relatively few genes remain heterogeneous after the imputation step, so the imputed *I*(*g*) identifies a much smaller number of genes than other methods, where an arbitrary cut-off in selected gene number is required.

## Conclusion

Traditional unsupervised clustering methods for single cell data are based on the, often implicit, assumption that cell types correspond to regions of high probability density in the joint gene expression distribution (Greulich et al. 2020). Although useful, this is not a specifically biological assumption (similar assumptions form the basis of numerous clustering algorithms in other different disciplines) and does not have a clear biological basis. Indeed, this assumption does not naturally accord with biological intuition that distinct cell populations should be ‘homogeneous’ and cells of the same type should be functionally interchangeable. Here, we have taken a different approach that seeks to encode biological intuition about population homogeneity in a mathematical formulation, drawing on tools from information theory.

We have proposed a formal measure of gene expression heterogeneity and shown that this measure captures biologically relevant heterogeneity arising from differential expression between established cell types. We formalised the additive decomposition of heterogeneity with respect to a given grouping of cells (in **Methods**), and tested the association between high inter-cluster heterogeneity and grouping by cell type. Finally, we used this measure as a basis of a biologically-motivated clustering procedure, which we call single-cell entropic clustering (scEC), which identifies homogeneous cell types. The scEC method is free of tunable hyper-parameters and its performance is comparable to a state-of-the-art clustering algorithm on a series of benchmark sc-Seq data sets, suggesting that this underlying biological basis is justified.

While our method represents a mathematically rigorous definition of heterogeneity, the stochasticity inherent in sc-Seq data does limit our resolution. For instance, nearly every gene is associated with some non-zero level of heterogeneity and, over thousands of genes, even low levels of stochastically-induced heterogeneity will accumulate (a version of the so-called ‘curse of dimensionality’). This limits our ability to directly interpret the genome-wide heterogeneity (e.g. the total heterogeneity of the technical control *Svensson* exceeds that of *Tian*, both absolute and per gene, although the trend reverses when excluding those genes with fewer than 100 total transcripts).

Aside from these technical limitations, our approach demonstrates the benefit of a biologically grounded, mathematically formal approach to understanding sc-Seq data. As these data sets grow in complexity and size, there will be a greater need for biological interpretability and mathematical rigour in analysis. We suggest information theory as the appropriate mathematical language for describing gene expression and its heterogeneity. Indeed, we anticipate that information theory will become an established part of the quantitative biologist’s toolkit.

## Methods

### Data collection

Count matrices for each data set were downloaded from their respective repositories (see **Availability of Data and Materials**). For the *Zheng* and *Tabula Muris* data sets, the count matrices of the various cell types/tissues were concatenated using the `Matrix` (v1.3-2) package in R (v4.0.3) (R Core Team 2020).

### Data pre-processing

For each data set, those genes with less than 100 total transcripts in total (i.e. across all cells) were excluded from further analysis. For the *Zheng* data set, FACS identities were taken from the repository metadata, with the CD4+ Helper T-cells, CD4+/CD25+ Regulatory T-cells, CD4+/CD45RA+/CD25- Näive T cells, CD4+/CD45RO+ Memory T-cells, CD8+ Cytotoxic T cells and CD8+/CD45RA+ Näive Cytotoxic T-cells merged into the single identity of T-cells.

### Normalisation

Since sc-Seq data is noisy, rather than working with the experimentally observed proportions of transcripts assigned to each cell, we instead adopt a Bayesian approach to estimate the cellular frequencies using the James-Stein estimator, as explained below, and use this estimator in all subsequent calculations.

We write 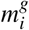 for the number of transcripts of gene *g* associated with cell *i* in a population of *N* cells. Let 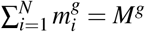 be the total number of transcripts of gene *g*, and 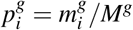 the fraction of transcripts of gene *g* expressed by cell *i*, for each 1 *≤ i ≤ N*. Thus, 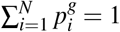 and we can think of 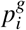 as the probability that a randomly selected transcript of gene *g* is associated with cell *i*. To take under-sampling into account, we adjust the observed probabilities 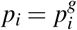 as follows (let us drop the superscript *g* for simplicity). We define *X* = *X*^*g*^ as the discrete random variable on the set *{*1, 2, … *N}* with probabilities

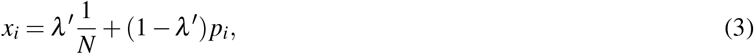

where *λ* ^*l*^ is the shrinkage intensity given by

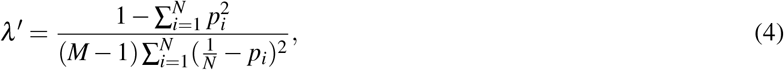

which is the James-Stein estimator, with the uniform distribution as the shrinkage target (Hausser & Strimmer 2009). Shrinkage approaches deal with under-sampling and so are able to correct for the substantial sparsity observed in single-cell RNA-sequencing data. The shrinkage probabilities *x*_*i*_ given in **Eqn. 3** are a compromise between the observed probabilities *p*_*i*_, which are unbiased but with a high variance, and the target uniform probabilities 1*/N*, which are biased but with low variance (Hausser & Strimmer 2009, Chan et al. 2017).

### Feature selection

The Shannon entropy of *X* is, by definition,

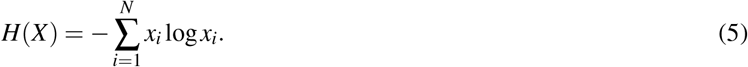

By convention, we assume that 0 *·* log 0 = 0. For the logarithm, we take base *e* for ease in differentiation (see later) rather than the usual base 2 (recall that log_*e*_(*x*) = log_2_(*e*) log_2_(*x*) for all *x >* 0), and thus *H*(*X*), and *I*(*g*) below, are measured in nats.

The Shannon entropy is a measure of the uncertainty in the outcomes of the random variable *X*. It has a minimum value of zero, when *x*_*i*_ = 1 for some *i* (e.g. if the gene *g* is expressed in only one cell of the population) and has a maximum value of log(*N*), when *x*_*i*_ = 1*/N* for all *i* (e.g. if the gene is uniformly expressed in the cell population). The entropy may therefore be considered as a measure of the *homogeneity* of expression of the gene *g* in the cell population profiled (note that the association of high entropy with homogeneity is counter to the usual intuition: it occurs because we are concerned with the entropy of the generative process of assigning transcripts to cells, rather than transcript count distribution in the cell population, as is more usual. A high entropy transcript assignment process gives rise to a sharp transcript distribution, hence its association with homogeneity). By contrast, the quantity log(*N*) *−H*(*X*) also ranges between zero and log(*N*), yet is minimised when the gene is homogeneously expressed and so is a simple measure of expression *heterogeneity*, which we will denote *I*(*g*). We can rewrite this as

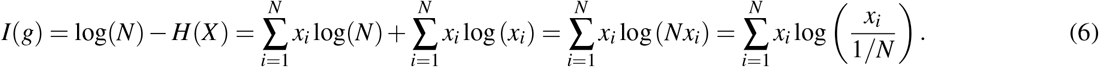

The Kullback-Leibler divergence (KLD), or relative entropy, is a measure of the information encoded in the difference between two probability distributions (Kullback & Leibler 1951). The relative entropy of a discrete probability distribution *p*_1_, …, *p*_*N*_ from a discrete probability distribution *q*_1_, …, *q*_*N*_ is, by definition,

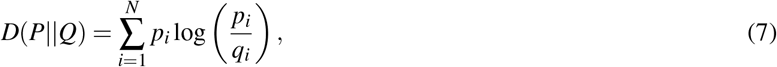

with the provision that *q*_*i*_ = 0 implies *p*_*i*_ = 0, and the convention that *·*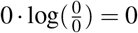. From this definition and **Eqn. 6**, it is clear that our measure of heterogeneity is simply the relative entropy of the observed expression distribution from the uniform distribution *U*. Thus,

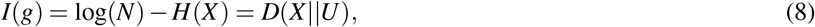

where *U* denotes to the uniform distribution on the set *{*1, 2, … *N}* with probabilities *u*_*i*_ = 1*/N* for all 1 *≤ i ≤ N*.

### Cluster-level heterogeneity

A crucial property of the relative entropy is that it is additively decomposable with respect to arbitrary groupings (Shorrocks 1980). Informally, this means that if we have a clustering of the cells into disjoint groups, then *I*(*g*) can be reconstructed from inter-cluster and intra-cluster heterogeneities. Next, we formalise this additive decomposition and give a self-contained derivation (cf. Theil (1967)).

Consider a clustering *S* = *{S*_1_, …, *S*_*C*_*}* that unambiguously assigns each cell in the sample into one of *C* non-intersecting sub-populations *S*_1_, …, *S*_*C*_ of sizes *N*_1_, …, *N*_*C*_. Note that 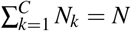, the total number of cells. Let *y*_*k*_ be the fraction of transcripts associated with cells in sub-population *S*_*k*_, adjusted by the shrinkage estimator. That is,

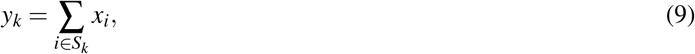

with *x*_*i*_ defined by **Eqn. 3**. This gives another discrete random variable *Y* with probability distribution *y*_1_, …, *y*_*C*_ on the set *{*1, 2, …*C}*. For each *k* = 1, …, *C*, we can also assess the heterogeneity of the sub-population *S*_*k*_ by considering the random variable *Z*_*k*_ with probability distribution *z*_*i*_ = *x*_*i*_*/y*_*k*_ on the set *i ∈ S*_*k*_. Note that

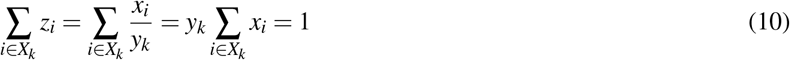

so *Z*_*k*_ is a random variable on the set (cluster) *S*_*k*_.

We may rewrite *I*(*g*) in terms of *Y* and *Z*_*k*_, as follows:

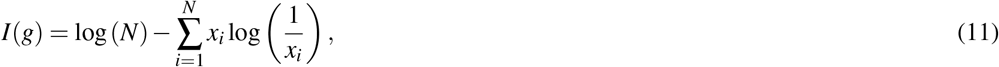

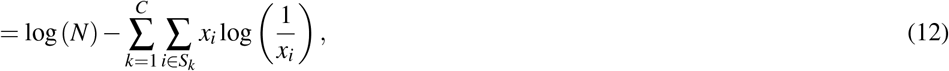

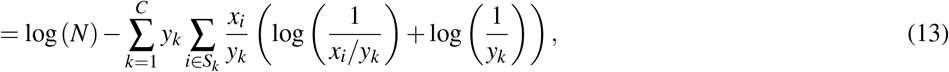

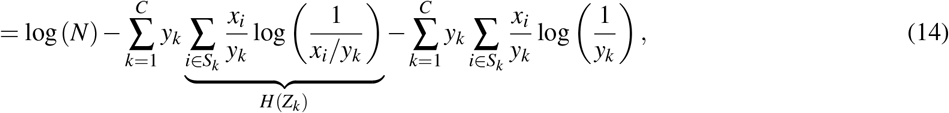

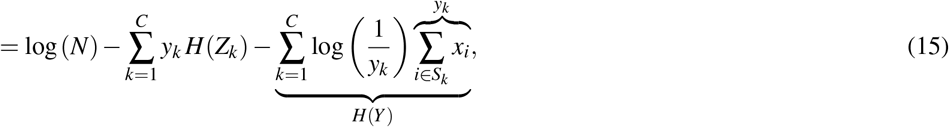

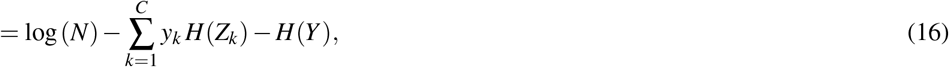

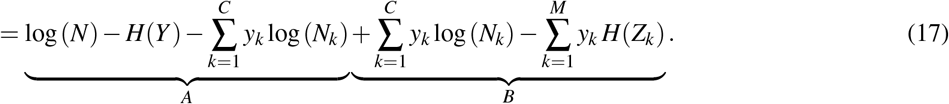

Expression *A* may be rewritten as:

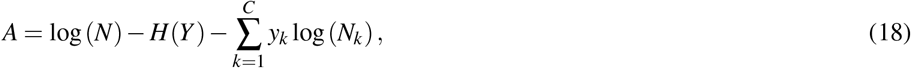

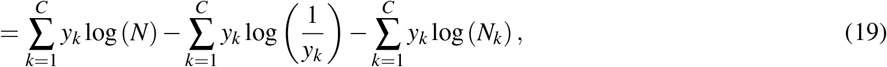

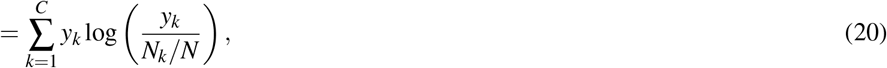

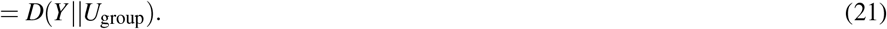

This is the relative entropy of *Y* from the uniform distribution *U*_group_ in which *p*_*k*_ = *N*_*k*_*/N* for *k* = 1, …, *C*. Since *y*_*k*_ is the proportion of transcripts assigned to the cluster *S*_*k*_, it measures the deviation from the assumption that the clusters are homogeneous in their expression of *g* (i.e. each cluster expresses *g* at the same level). Since it is a measure of the extent to which the population deviates from homogeneity between clusters, we will term this contribution the *inter-cluster heterogeneity* of *g* with respect to the clustering *S*, denoted *H*_*S*_. Informally, it is a measure of the extent to which the gene *g* is differentially expressed between clusters.

Expression *B* may be rewritten as:

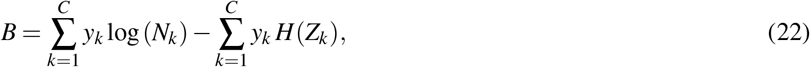

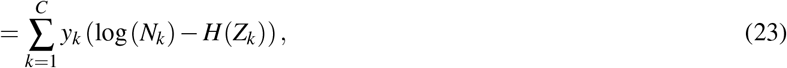

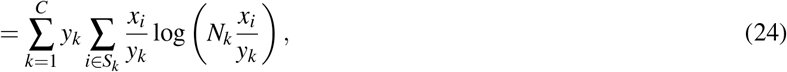

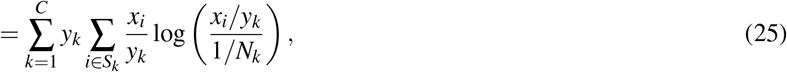

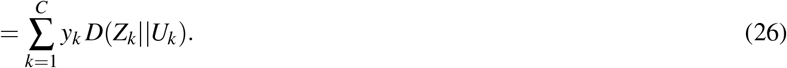

This is the weighted sum of the relative entropies of the empirical distributions *Z*_*k*_ (i.e. the observed gene expression distribution in group *S*_*k*_) from the uniform distribution *U*_*k*_ on *S*_*k*_ (in which *p*_*i*_ = 1*/N*_*k*_ for each *i ∈ S*_*k*_). It is the deviation from the assumption that the population consists of a mixture of homogeneous sub-populations according to the clustering *S* (where the expectation is taken with respect to the probability measure provided by *Y*). Since it is a measure of the expected extent to which the proposed sub-populations deviate from homogeneity within clusters, we will term this contribution the *intra-cluster heterogeneity* of *g* with respect to *S*, denoted *h*_*S*_(*g*).

Taken together, these results show that *I*(*g*) can be decomposed into two well-defined parts that encode local and global properties of the expression distribution of *g* with respect to a given clustering *S* = *{S*_1_, …, *S*_*C*_*}*:

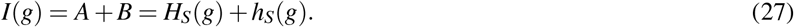

The relative entropy is always non-negative and hence so are *H*_*S*_(*g*) and *h*_*S*_(*g*) for any *S*. Thus, both quantities range from zero to *I*(*g*). If *S* places one cell in each cluster (i.e. *C* = *N*) then *Z*_*k*_ = *U*_*k*_ for all *k* and thus *h*_*S*_(*g*) = 0. Conversely, if *S* places all cells in one cluster (i.e. *C* = 1) then *Y* = *U*_*S*_ and thus *H*_*S*_(*g*) = 0. In this case, the population heterogeneity is equivalent to the intra-cluster heterogeneity of the trivial clustering.

### Unsupervised clustering

The above measures of heterogeneity can be easily extended from one gene to many. In this case, the homogeneous null model is obtained by assuming that each gene is expressed homogeneously and independently. Because entropy from independent sources is additive (Cover & Thomas 2012), the total heterogeneity of a single-cell RNA-sequencing data set under a given clustering is given by the sum:

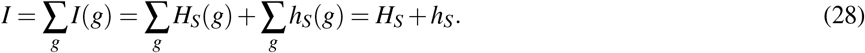

Here *H*_*S*_ = ∑_*g*_ *H*_*S*_(*g*) and *h*_*S*_ = ∑_*g*_ *h*_*S*_(*g*) represent the total inter-cluster and intra-cluster heterogeneity of the data with respect to a clustering *S* = *{S*_1_, …, *S*_*C*_*}*.

Naturally, we want to identify a clustering (e.g. an assignment of cells to types) that produces maximally homogeneous sub-groups and thereby explains most of the expression heterogeneity for most genes in terms of inter-cluster heterogeneity. Mathematically, this means we want to find a clustering *S* that maximises *H*_*S*_ or, equivalently, minimises *h*_*S*_. A brute force approach is not feasible, as the number of partitions of a set with *n* elements into two or more subsets grows exponentially with *n*.

To approach the problem we therefore converted the problem from one of assigning discrete identities (i.e., a hard clustering problem) to one of assigning continuous identities (i.e., a fuzzy clustering problem). The advantage of this approach is that it defines the problem in terms of a continuous, differentiable objective function which can be approached with more efficient optimisation routines.

#### Fuzzy clustering

We adopt a fuzzy conception of clustering in which cells are assigned to *C* (possibly) overlapping, fuzzy clusters, *S*_1_, …, *S*_*C*_. Each cell *i* has *C* corresponding membership functions *μ*_*ik*_ *∈* [0, 1] for *k* = 1, 2, …, *C*, which assess the probability that cell *i* belongs to cluster *S*_*k*_. Thus,

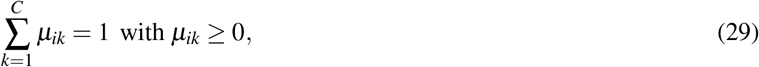

Our information-theoretic framework can be adapted to this fuzzy setting. Population heterogeneity, *I*(*g*), is independent of the chosen clustering, so it remains unaffected by the adoption of fuzzy clusters, i.e. the total gene expression heterogeneity is unaffected by the choice of discrete or fuzzy cluster memberships. For the calculations of inter-cluster and intra-cluster heterogeneities, we extend the discrete random variables *Y*^*g*^ and 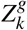 to the fuzzy setting, where *y*^*g*^ now measures the expression distribution of the gene *g* across the *C* fuzzy clusters, and 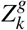 measures the expression distribution of the gene *g* within fuzzy cluster *S*_*k*_, as follows.

We define the discrete random variable *Y*^*g*^ on the set of fuzzy clusters *S* = *{S*_1_, …, *S*_*C*_*}* with probabilities 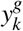 given by

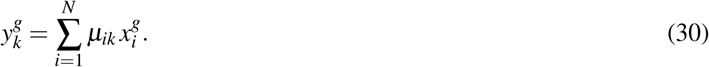

We also define, for each *S*_*k*_ *∈ S*, a discrete random variable 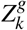 on the set of cells, *i* = 1, …, *N*, with probabilities 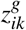 given by,

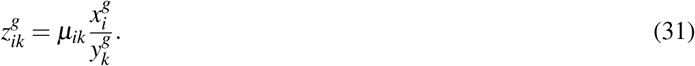

We can rewrite *I*(*g*) (which is independent of the clustering) in terms of *Y*^*g*^ and 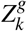 (which depend on the clustering) as

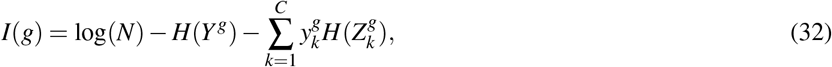

where,

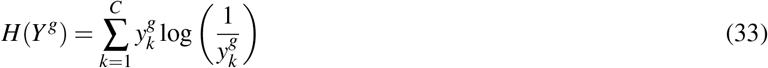

And

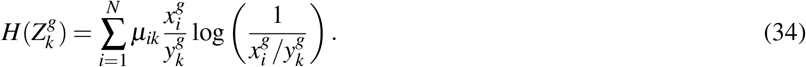

Note that **Eqn. 33** is the entropy of the random variable *Y*^*g*^, however 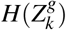 is not quite the entropy of 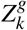 (there is a missing 1*/μ*_*ik*_ factor inside the logarithm).

From **Eqn. 32**, we can obtain a second decomposition,

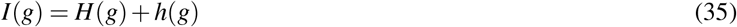

Where

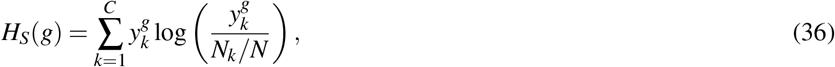

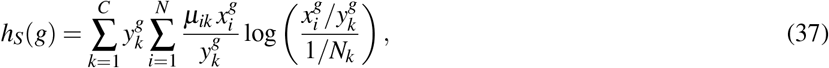

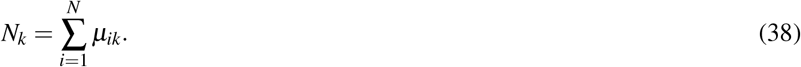

Here *H*(*g*) and *h*(*g*) represent the inter-cluster heterogeneity and intra-cluster heterogeneity of gene *g* with respect to the fuzzy clustering *S*. **Eqn. 36** is also the KLD of the distribution of *Y*^*g*^ from the uniform distribution *U*_group_ given by *u*_*k*_ = *N*_*k*_*/N* for *k* = 1, 2, …, *C* (note that 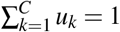. Similarly, **Eqn. 37** is the KLD of the distribution of *Z*^*g*^ from the uniform distribution *U*_*k*_ given by *p*_*i*_ = *μ*_*ik*_*/N*_*k*_ for *i* = 1, 2, …, *N*.

Either *H*_*S*_ = ∑_*g*_ *H*_*S*_(*g*) or *h*_*S*_ = ∑_*g*_ *h*_*S*_(*g*) can serve as our objective function, *f*, for optimisation: optimising either quantity simultaneously identifies the clustering with the greatest heterogeneity between clusters (i.e. the greatest differential gene expression between clusters) and the least heterogeneity within clusters (i.e. in which the cells within each cluster are closest to being interchangeable with respect to their patterns of gene expression).

Choosing inter-cluster heterogeneity *H*_*S*_, we solve the optimisation problem

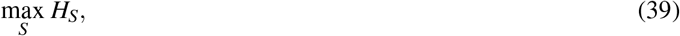

Where

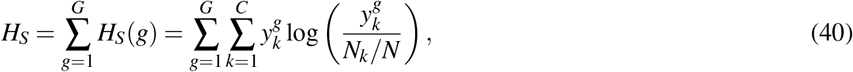

using the limited-memory, box constrained BFGS (*L-BFGS-B*) optimisation algorithm from the `Python3` (v3.8.2) `package SciPy` (v1.5.3) (Byrd et al. 1995, Zhu et al. 1997, Van Rossum & Drake 2009, Virtanen et al. 2020).

The *L-BFGS-B* algorithm is an efficient non-linear local optimisation method developed for solving large, dense problems such as this (Zhu et al. 1997)) but its speed and the quality of the solution produced depends on the availability of an explicit formulation of the gradient of the objective function. Therefore, we derive the required gradient by partially differentiating the total inter-cluster heterogeneity, *H*_*S*_, with respect to fuzzy cluster membership, i.e. the *N · C* membership functions *μ*_*rs*_ (1 *≤ r ≤ N*, 1 *≤ s ≤ C*).

Beginning by differentiating individual elements of *f* with respect to *μ*_*rs*_, from **Eqn. 30** and **38**, we have

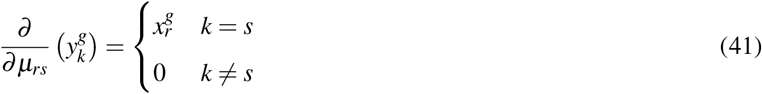

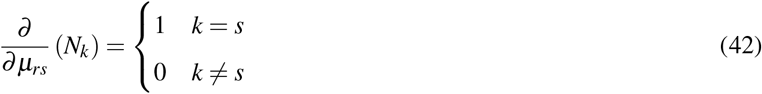

Using the product, chain and quotient rules, we can differentiate the *g*-summand, call it *f* ^*g*^ (= *H*_*S*_(*g*)), of the objective function *f* :

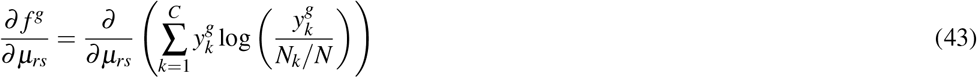

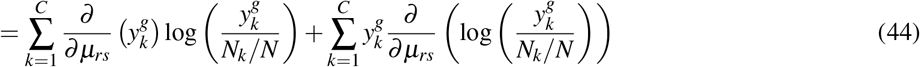

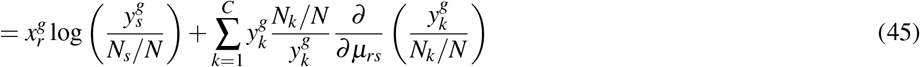

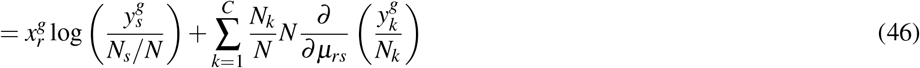

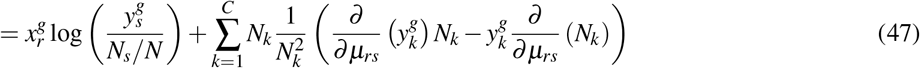

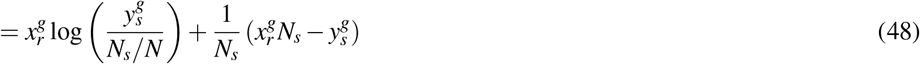

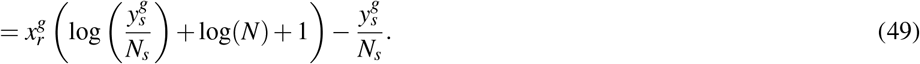

The membership functions *μ*_*rs*_ are not independent, since 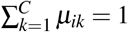 for all *i*, by **Eqn. 29**. To incorporate this constraint, and the constraint *μ*_*ik*_ *≥* 0 for all *i, k*, we introduce variables *w*_*ik*_ given by

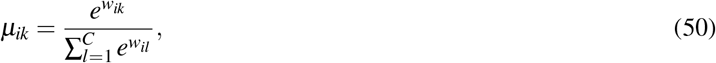

and make the objective function *f* a function of the *w*_*ik*_. By the chain rule,

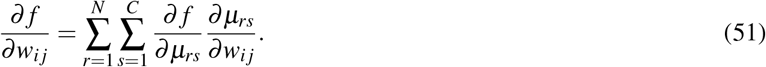

Determining 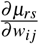, let us write 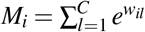 ; then, **Eqn. 50** can be written as 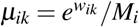. By the quotient rule,

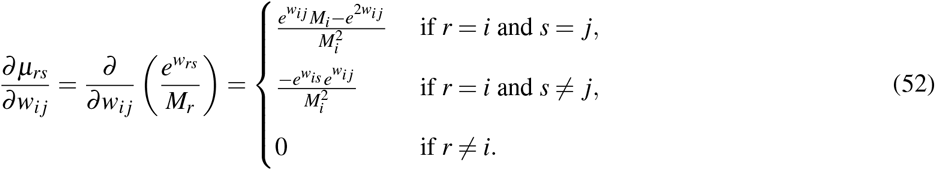

#### Numerical optimisation

The objective function *H*_*S*_, as formulated in **Eqn. 40**, was optimised with respect to cluster memberships, *μ*_*ik*_ using the *L-BFGS-B* optimisation algorithm from the `Python3` (v3.8.2) `package SciPy` (v1.5.3) (Byrd et al. 1995, Van Rossum & Drake 2009, Virtanen et al. 2020), with random initial starting points (initial cluster memberships were defined by random sampling of a uniform distribution centred on zero). The gradient function, as formulated in **Eqn. 51**, was supplied to the optimisation routine.

Because initial conditions are randomly generated, and the *L-BFGS-B* algorithm only identifies local optima, the specific clustering found by the algorithm may depend on the choice of initial vector (Byrd et al. 1995, Zhu et al. 1997). To make the implementation robust a multi-start process is adopted, in which the optimisation is repeated multiple times with different initial vectors, and the clustering *S* with the greatest inter-cluster heterogeneity *H*_*S*_ is chosen.

#### Permutation testing

Annotations for the *Tian, Zheng* and *Stumpf* data were taken from the respective repositories’ metadata. Random clusterings were generated by randomly permuting the established annotations 10,000 times each. Comparison of true and shuffled annotations was formulated as an exact one-side hypothesis test, generating exact *p*-values (Fisher 2017). We control the false discovery rate arising from multiple testing for each data set using the R function *p*.*adjust* (Benjamini & Hochberg 1995).

#### Clustering comparison

We compare our clustering results to two standard clustering methods: `Seurat` (v3) clustering (Louvain community detection), and UMAP projection, both with default parameters, with the exception of the resolution, as described in Stuart et al. (2019) (Blondel et al. 2008, McInnes et al. 2018). The Adjusted Rand Index (ARI) was calculated for each data set using the associated function in the R package `mclust` (v5.4.7) (Rand 1971, Scrucca et al. 2016).

##### Semi-supervised classification

Consider a pair of count matrices, *T* = *{T*_*re f*_, *T*_*test*_*}*, where one count matrix, which we term the reference, *T*_*re f*_, has a known cluster structure (and therefore each cell has a unique, discrete cell type classification). The goal of reference mapping is to classify cells in the test matrix, *T*_*test*_, based on cellular classifications of the reference matrix.

We normalise each count matrix separately as described in **Normalisation**, using the James-Stein-type estimator so that 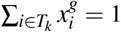 (Hausser & Strimmer 2009). We assume that the same set of genes are profiled in *T*_*re f*_ and *T*_*test*_ and derive the normalised combined count matrix, *X*_*mix*_, via the weighted concatenation:

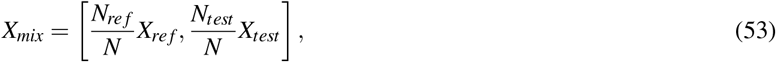

where *N*_*re f*_ is the number of cells in the reference data set, *N*_*test*_ is the number of cells in the unclassified data set, and *N* is the total number of cells across both data sets.

As in the unsupervised setting, we cluster the combined data set, *X*_*mix*_, by maximising *H*_*S*_ of the combined data set with respect to a fuzzy clustering *S*, see **Eqn. 56**. However, unlike in the unsupervised setting, a subset of cellular cluster memberships are known *a priori*. Let *R* = *R*_1_, …, *R*_*C*_ be the discrete clustering of the reference data set, with cluster sizes *n*_1_, …, *n*_*C*_, where 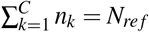. As the cluster identities of cells originating from the reference data set are fixed, each cluster in the reference data set constitutes a subset of a cluster in the mixed data set, *R*_*k*_ *∈ S*_*k*_, where *S* = *S*_1_, …, *S*_*C*_ is the clustering of the mixed data set. Note that number of clusters in the reference and in the combined data set is the same, i.e. *C*.

Based on the co-mixing of known and unknown cellular identities, 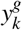 and *N*_*k*_ of the combined data set *X*_*mix*_ are given by

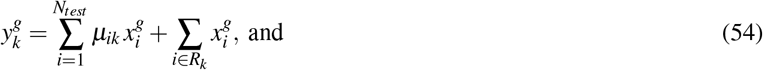

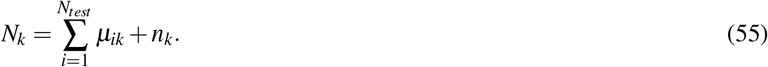

Based on these formulations of 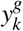 and *N*_*k*_, the definition of *H*_*S*_ follows as before,

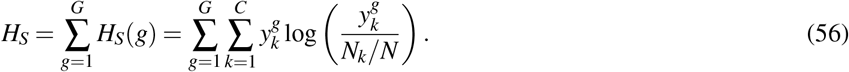

Similarly, the derivative of *H*_*S*_ follows as before, except that the membership function, *μ*_*rs*_, is only defined with respect to cells of the test data set, 1 *≤ r ≤ N*_*test*_ *&* 1 *≤ i ≤ N*_*test*_,

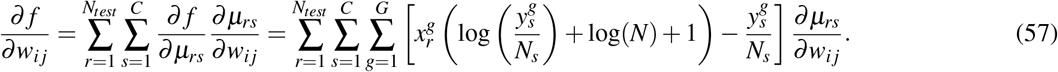

Numerical optimisation follows as in the unsupervised setting, using the *L-BFGS-B* algorithm, except that the known cellular identities of the reference mean that only a single run of the algorithm is required, i.e. no multiple starts.

### Imputation

We rely on a cell-by-cell adjacency matrix for imputation, encoding which cells have similar gene expression profiles. Specifically, we use the shared-nearest-neighbours matrix produced by `Seurat`, using default settings (Stuart et al. 2019).

We begin with the expression matrix *X*, keeping only the top *G* genes by *I*(*g*) (default is 500). Then, briefly, the `Seurat` method follows the following steps: data is scaled and normalised using a variance stabilising transform; the transformed data then undergoes principal components, and a cell-cell Euclidean distance matrix is calculated on the first 10 principal components. Next, a *k*-nearest-neighbours graph is constructed, where *k* is by default 20; a shared nearest-neighbours graph is constructed by taking the Jaccard Index of the overlap in neighbourhoods of each pair of cells (Jaccard 1912).

From this adjacency matrix, *A*, we compute the imputed data via a one-hop smoothing operator:

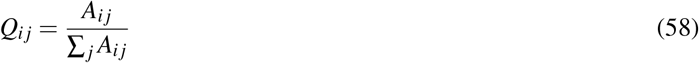

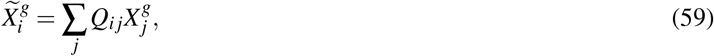

where *Q* is the row-normalised stochastic matrix, i.e. 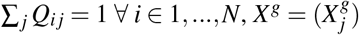 is the *g*-column of *X*, and 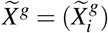 are the imputed values for gene *g* (Ortega 2022). The imputed expression value of gene *g* in cell *i* is a weighted mean of observed expression values in cell *i* and its neighbourhood in *A*. This is an example of a data diffusion imputation, first applied to Sc-seq data in Van Dijk et al. (2018).

The vector 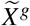 can then be used in place of *X*^g^ in any of the above described work. In the main text, we select the top 500 genes by *I*(*g*) based on 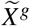, before clustering on 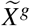. Accordingly, use of the imputed expression matrix requires two rounds of feature selection: first in the construction of the adjacency matrix, *A*, then for the use of 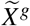 in clustering.

For the benchmarking of the imputation, we recapitulated the set of data sets and methods tested in Tian et al. (2019).

However, we were unable to rerun RCA (Li et al. 2017).

## Declarations

## Authors’ contributions

Conceptualization, MJC, RSG, BDM; Software, MJC; Investigation, MJC, JF, RJSG and BDM; Writing – Original Draft, MJC, RSG and BDM; Writing – Review & Editing, MJC, JF, RSG and BDM; Visualization, MJC; Supervision, RSG and BDM. All authors have seen and approved the manuscript.

## Acknowledgements

We thank P. Stumpf, J. Egan and B. Kitching-Morley for helpful discussions.

## Availability of data and materials

The *Svensson* data was downloaded as file “svensson chromium control.h5a” from https://data.caltech.edu/records/1264

The *Tian* data sets were downloaded from https://github.com/LuyiTian/sc_mixology

The *Zheng* data was downloaded as set of files “Gene / cell matrix (filtered)” in section “Single Cell 3’ Paper: Zheng et al.

2017” from https://support.10xgenomics.com/single-cell-gene-expression/datasets

The *Stumpf* data was downloaded as file “RData.zip” from https://data.mendeley.com/datasets/csvm3kpkxd/1?fbclid=IwAR3j_hvq5Zt0cdxBBM72Fqr8zXwVC6XRpkY2JtVwbcj1gzyNLMVsdoFYWgQ

The *Tabula Muris* data was downloaded as set of files “Single-cell RNA-seq data from microfluidic emulsion (v2)” from

https://tabula-muris.ds.czbiohub.org

## Code availability

An R package for the implementation of the described methods is available on github https://github.com/mjcasy/scEC

## Ethics approval and consent to participate

Not applicable.

---

**Supplementary Figure 1.**
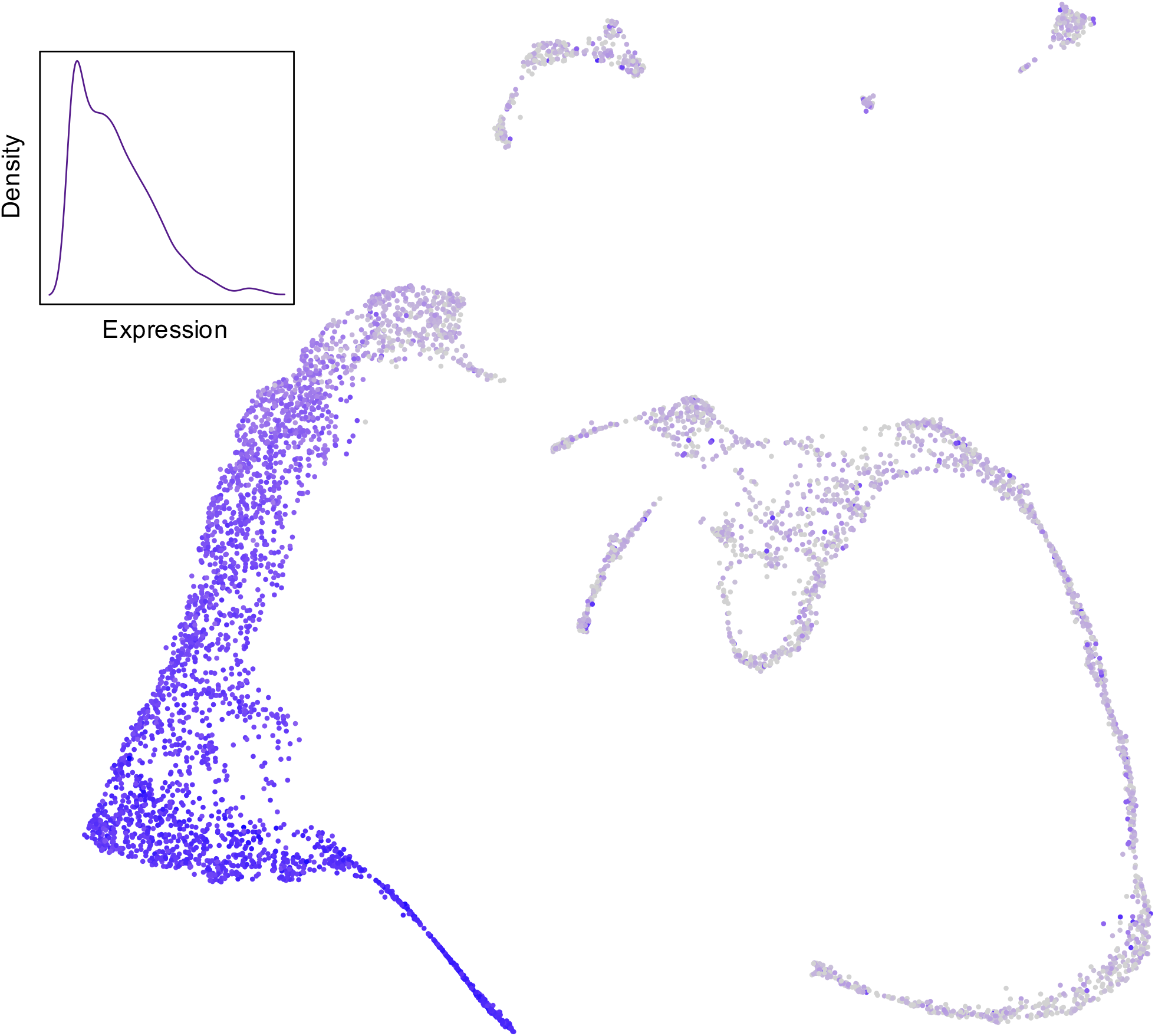
Expression of *Hbb-bs* in Stumpf et al. (2020) data set. *Hbb-bs* is identified by information-theoretic feature selection – based on *I*(*g*) – but not by scran (Lun et al. 2016); *Hbb-bs* has a complex distribution, with at least two modes of expression in erythroblasts (inset shows probability density, y-axis, of transcript counts, x-axis, in erythrocyte lineage), with a high level of expression in mature erythrocytes and some non-zero level of (possible technically-induced) expression across less developed erythroblasts and other cell types. The non-parametric nature of *I*(*g*) means it can robustly identify informative genes, even in the presence of multimodal expression patterns.

**Supplementary Figure 2.**
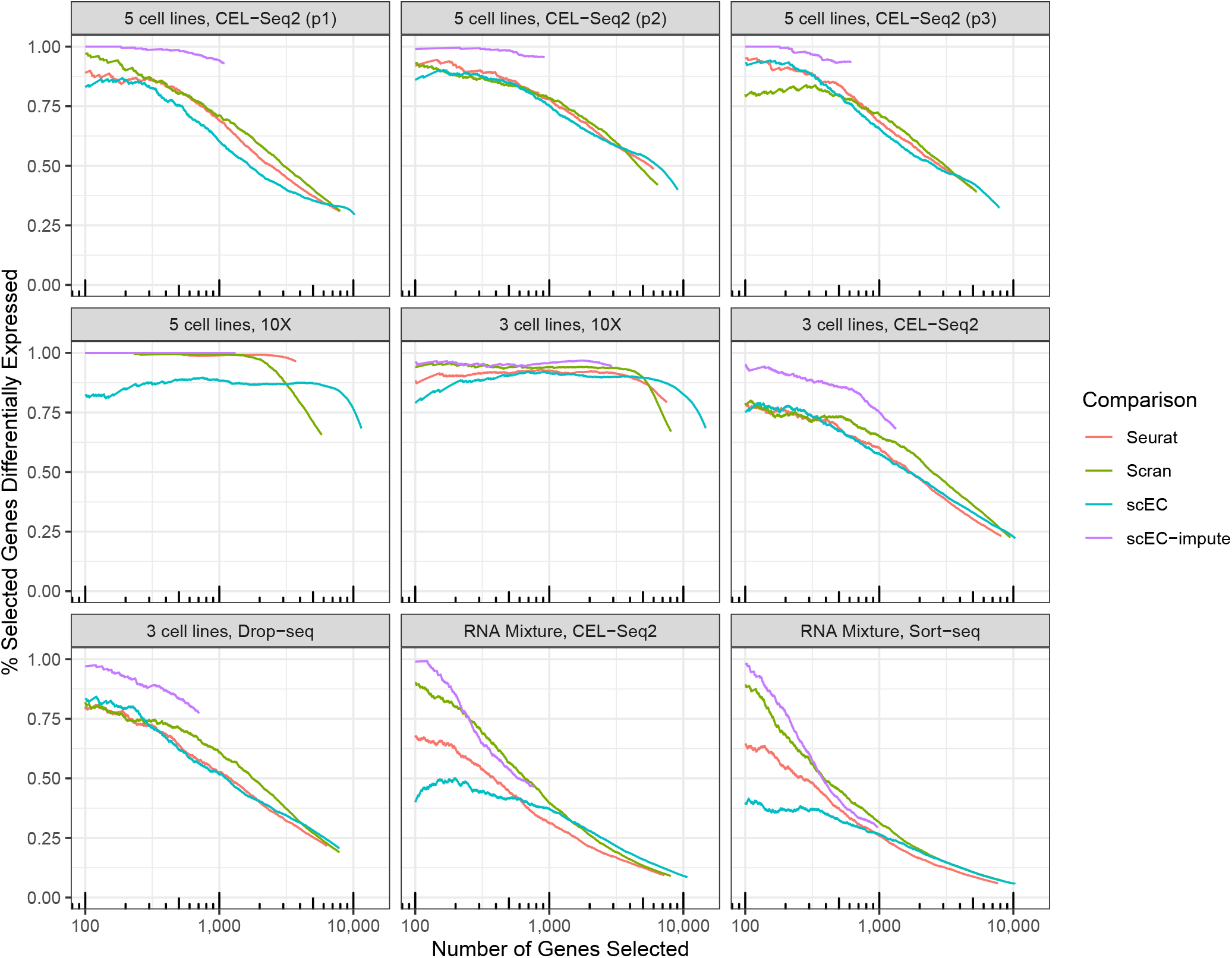
Feature selection benchmarking of scEC and scEC-impute. he percent of the top *N* genes by different feature selection metrics that are differentially expressed. Data is Sc-seq from three or five cancerous cell lines, or a mixture of RNA from said cell lines, sequenced by specified platform (Tian et al. 2019). In each case, scEC-impute has the greatest fidelity in selecting differentially expressed genes without access to cluster labels, with the exception of the RNA mixture data sets where it draws equal with the other methods after several hundred genes have been selected. Number of differentially expressed genes identified in each study by Wilcox test of each cell line against remaining, false discovery rate corrected *p*-value *<* 0.05: 5 cell lines, CEL-Seq2 (p1) 3695; 5 cell lines, CEL-Seq2 (p2) 4523; 5 cell lines, CEL-Seq2 (p3) 3390; 5 cell lines, 10X 7851; 3 cell lines, 10X 10195; 3 cell lines, CEL-Seq2 2712; 3 cell lines, Drop-seq 2005; RNA Mixture, CEL-Seq2 947; RNA Mixture, Sort-seq 667.

**Supplementary Table.** Contingency table between cell annotations provided in (Stumpf et al. 2020) and scEC labels. Note that while the number of clusters for scEC was fixed at 14, only 13 clusters were realised; generally, the number of clusters returned will match the number specified, except in cases where any further splits in clusters provide only a minor increase in inter-cluster heterogeneity, so are harder to find by numerical optimisation.

|  |  |  |  |  |  |  |  |  |  |  |  |  |  |  |
| --- | --- | --- | --- | --- | --- | --- | --- | --- | --- | --- | --- | --- | --- | --- |
| Erythroblasts | 2436 | 171 | 0 | 0 | 0 | 1 | 0 | 0 | 1 | 0 | 233 | 0 | 3 | 0 |
| Myeloblasts | 39 | 4 | 313 | 0 | 0 | 0 | 0 | 0 | 0 | 0 | 0 | 245 | 2 | 0 |
| Pro-B | 38 | 0 | 0 | 187 | 0 | 0 | 0 | 0 | 0 | 0 | 0 | 0 | 0 | 0 |
| Pre-B | 0 | 0 | 0 | 58 | 0 | 0 | 0 | 0 | 2 | 0 | 0 | 2 | 0 | 0 |
| Monocytes | 0 | 0 | 0 | 23 | 165 | 0 | 0 | 0 | 4 | 2 | 0 | 23 | 0 | 0 |
| Basophils | 0 | 0 | 0 | 0 | 0 | 130 | 0 | 0 | 0 | 0 | 2 | 1 | 0 | 0 |
| Pericytes | 0 | 0 | 0 | 0 | 0 | 0 | 110 | 0 | 0 | 2 | 0 | 0 | 0 | 0 |
| Megakaryocytes | 0 | 0 | 0 | 0 | 0 | 0 | 0 | 40 | 0 | 0 | 21 | 0 | 0 | 0 |
| T-NK | 3 | 0 | 0 | 0 | 0 | 0 | 0 | 0 | 56 | 0 | 0 | 1 | 0 | 0 |
| Endothelial Cells | 0 | 0 | 0 | 0 | 1 | 0 | 4 | 0 | 0 | 41 | 4 | 0 | 0 | 0 |
| HSPCs | 49 | 4 | 3 | 5 | 1 | 7 | 0 | 0 | 2 | 0 | 182 | 50 | 0 | 0 |
| Monoblasts | 7 | 0 | 16 | 7 | 66 | 0 | 1 | 0 | 0 | 0 | 3 | 200 | 0 | 0 |
| Neutrophils | 0 | 0 | 0 | 1 | 1 | 1 | 0 | 0 | 1 | 0 | 0 | 8 | 255 | 0 |
| Myelocytes | 2 | 0 | 4 | 3 | 0 | 0 | 0 | 0 | 0 | 0 | 0 | 93 | 164 | 0 |

